# Anisotropic dense collagen hydrogels possessing two ranges of porosity to create the adequate microenvironment for muscle bundles: a step towards skeletal muscle modeling

**DOI:** 10.1101/2022.06.19.496716

**Authors:** Marie Camman, Pierre Joanne, Julie Brun, Alba Marcellan, Julien Dumont, Onnik Agbulut, Christophe Hélary

## Abstract

Despite the crucial role of the extracellular matrix (ECM) in the organotypic organization and function of skeletal muscles, most 3D models do not mimic its specific characteristics, namely its biochemical composition, stiffness, anisotropy, and porosity. Here, a novel 3D *in vitro* model of muscle extracellular matrix was developed to differentiate myogenic cells (C2C12 line) into myotubes and reproduce their natural cell/cell and cell/matrix interactions. An anisotropic hydrogel mimicking the perimysium was obtained thanks to unidirectional 3D printing of dense collagen with aligned collagen fibrils. The space between the different layers was tuned to generate an intrinsic porosity (100 µm) suitable for nutrient and oxygen diffusion. By modulating the gelling conditions, the mechanical properties of the construct reached those measured in the physiological muscle ECM. The addition of large channels (600 µm) by molding permitted to create a second range of porosity suitable for cell colonization without altering the physical properties of the hydrogel. C2C12 cells embedded in Matrigel®, seeded within the channels, organized in 3D, and differentiated into multinucleated mature myotubes. This organization reproduced the global muscular bundles, *i*.*e*., the endomysium encompassing myotubes. These results show that porous and anisotropic dense collagen hydrogels colonized with myoblasts are promising biomaterials to model skeletal muscle.

**Table of contents:** 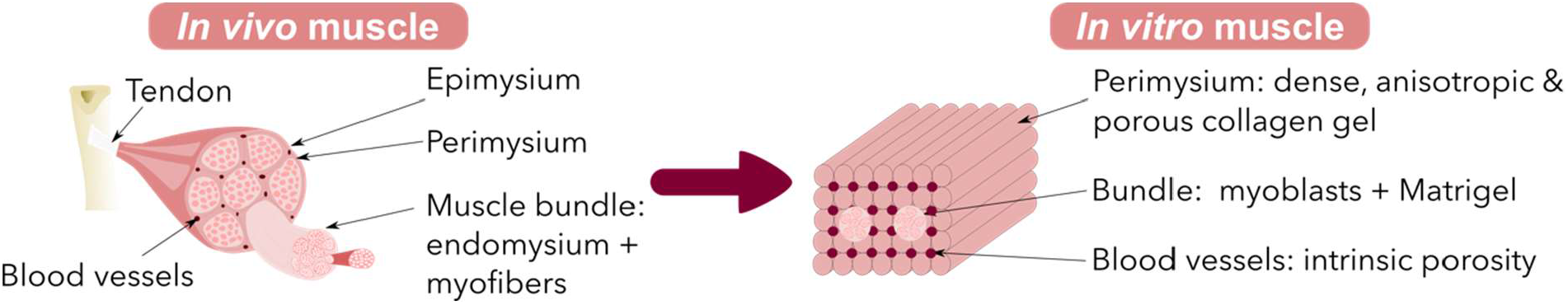

**Highlights:**

- A novel extracellular matrix-like hydrogel increases the physiological relevance of the skeletal muscle model.
- Porous and anisotropic dense collagen hydrogels mimic the muscle ECM physical properties.
- Unidirectional printing of dense collagen creates a porous and anisotropic scaffold in a single step.
- Anisotropic dense collagen hydrogels promote C2C12 differentiation into myotubes and their 3D organization.

## 1. Introduction

Skeletal muscles are involved in the body locomotion thanks to their highly organized tridimensional structure. The epimysium is a connective tissue surrounding the whole skeletal muscle. The perimysium, consisting of aligned collagen I fibers (anisotropy), separates each muscle bundle of myofibers [1]. These myofibers are highly aligned and densely packed within a loose connective tissue called endomysium. This latter resembles a basal membrane composed of collagen IV, fibronectin, and laminin [2]. This hierarchical organization is fundamental for muscle contraction and the integrity of the neuromuscular junction [3].

Because of its high complexity, the first *in vitro* models of skeletal muscle were limited to monolayers of myoblasts cultured on a petri dish coated with Matrigel® [4–6]. Matrigel reproduced the endomysium’s biochemical cues and helped myoblasts grow and differentiate. However, these 2D models did not copy the 3D muscle organization. 3D *in vitro* microtissues overcome these limitations by reproducing the cellular microenvironment. The most popular 3D models rely on encapsulating muscle cells within soft hydrogels made of fibrin, collagen, or gelatin between anchors [7–9]. Myoblasts align along the axis of the hydrogel due to cell contraction and create a displacement of the anchors. The primary readout of such a model is contractility. In these systems, the properties (stiffness and topography) of the muscular extracellular matrix, which are crucial in the muscle function for the force transmission, are not considered. In addition, hydrogels are not stable, and a necrotic core appears due to the absence of porosity.

The ideal biomaterial mimicking the physiological extracellular matrix (perimysium) surrounding the muscle bundles should be composed of 2-5% collagen I, be anisotropic and have high mechanical properties (E=12 kPa). In addition, this ECM should be porous to ensure nutrient diffusion [10] and allow myoblast survival. Most of the current models used synthetic or natural polymers but rarely collagen. This choice can be explained by the multiple drawbacks of collagen hydrogels when made from low concentrated solutions (0.5% wt): poor mechanical properties and shrinkage when colonized with cells [11].

Anisotropy of the muscle ECM is crucial to guide myofibers and allows their alignment *in vivo*. Mimicking the topography of the perimysium *in vitro* allows the control of the organotypic organization and the cell phenotype. Collagen anisotropy can be obtained by electrospinning or under a high magnetic field [12–15]. Besides, anisotropic collagen threads can be produced by extruding dense collagen solutions within a gelling bath made of PBS 5X [16] or other buffers [17]. Anisotropic layers or gels are obtained by 3D printing [18–20]. These strategies create oriented collagen fibrils (anisotropy) suitable for cell culture. However, they are not biomimetic 3D models when cells are seeded on top of the construct, *i*.*e*., not organized in densely packed bundles within the threads [21,22].

A significant limitation of 3D models is the slow diffusion of nutrients and oxygen, leading to the formation of a necrotic core [23]. Hence, porosity is an important feature to consider for cell survival maintenance. Several strategies, such as porogen leaching or self-assembly, have increased nutrient diffusion but require solvents or modified collagen. Additionally, pore geometry is barely controllable [24,25]. 3D printing can create an intrinsic porosity by stacking round filaments [26–28]. However, the 90° switch between layers disturbs anisotropy and the channel shape, limiting its utilization.

Other techniques, such as needle or suture thread molding, generate large and elongated channels suitable for cell colonization and differentiation into myotubes (>500 µm) [26,29–31]. This kind of porosity is challenging to obtain by 3D printing due to the channel collapsing. Unfortunately, it is also difficult to multiply the number of needles to dedicate one fraction for nutrient diffusion beside channels for cell colonization.

Hence, an artificial muscle ECM with the appropriate physicochemical cues and good mechanical properties has not been described yet. In addition, collagen biomaterials presenting two sets of porosity (for cell colonization and perfusion) have not been designed either.

In this study, we aim to design a novel *in vitro* 3D model mimicking the organization and the structure of the skeletal muscle, *i*.*e*., muscle bundles consisting of myotubes and loose extracellular matrix surrounded by the orientated dense collagen matrix. For this purpose, the key features of the muscle ECM (biochemical cues, stiffness, porosity, anisotropy) will be set within a single hydrogel using 3D printing. Then C2C12 cells, used as muscle cell models, will be seeded in 3D in Matrigel® to form a bundle inside the large channels of 3D printed hydrogels. Their differentiation into myotubes will thus be analyzed.

## 2. Materials and Methods

### 2.1. Collagen extraction and purification

As previously described, type I collagen was extracted and purified from rat tail tendons [32]. Briefly, rat tails were rinsed with ethanol 70% and cut into small pieces of 1 cm to extract tendons. Tendons were solubilized in 500 mM acetic acid. After precipitation with 0.7 M NaCl, centrifugation, and dialysis, collagen purity was observed after SDS-PAGE electrophoresis, and its concentration was estimated by hydroxyproline titration [32]. After an evaporation step in a safety cabinet for several days, collagen solutions concentrated at 30 mg.mL^-1^ (3%wt) in acetic acid were obtained. Finally, collagen solutions were stored at 4°C before utilization.

### 2.2. Collagen 3D printing

3D printing was performed using a home-built 3D printer. The concentrated collagen solution was poured into a 1 mL syringe (Terumo) with a flat bottom 23G needle (inner diameter 330 µm). A unique layer of 10 mm x 5 mm x 0.33 mm was designed using the AutoDesk Fusion 360 software. The 3D file (.stl) was then sliced with Repetier software with a rectilinear pattern to obtain unidirectional lines inside the square. This layer was printed and repeated at different z 5 times to form a 5-layered collagen construct with all filaments oriented in the same direction. The extrusion speed was optimal at 2 mm.s^-1^ (extrusion rate 1 µl.s^-1^). The filling was set to 100% to ensure cohesiveness between every filament but the interlayer gap varied between 300 and 500 µm to tune the hydrogel porosity. Anisotropy was induced by collagen shearing during extrusion, and this anisotropy was maintained by printing into a PBS 5X bath. The negative control was printed in air and gelled with ammonia vapors. The printing process lasted less than 10 minutes (1 min 30 per layer).

### 2.3. Collagen gelling and fibrillogenesis after 3D printing

Different strategies were tested to extend the collagen gelling and maintain the anisotropy while setting high mechanical properties. After the printing step inside PBS 5X, hydrogels were left in a large amount of PBS 5X to continue the initial gelling at pH 7.4 with high ionic strength (from 30 min to 7 days). At the end of this period, some collagen constructs were exposed to ammonia vapors for 24 hours to increase gelation For this purpose, an ammonium hydroxide solution (30% Carlo Erba) was poured into a glass beaker and placed in a desiccator. Constructs only gelled by ammonia vapors were printed in air and directly placed inside the desiccator for 1 day. After ammonia vapors, collagen constructs were rinsed several times in PBS 1X baths until a neutral pH was reached (at least 24 hours). Gelling conditions are presented in Table 1. Casted gels with the same gelling conditions were carried out as negative controls.

**Table 1.** Printing conditions and gelation process.

|  | Initial gelling | Extended gelling |  |
| --- | --- | --- | --- |
| Gelling and Fibrillogenesis Conditions | Printing bath (PBS 5X) | Time in PBS 5X | Time under ammonia vapors |
| 1 day NH<sub>3</sub> | No | - | 1 day |
| 30 min PBS/1 day NH<sub>3</sub> | Yes | 30 min | 1 day |
| 1 day PBS/1 day NH<sub>3</sub> | Yes | 1 day | 1 day |
| 6 days PBS/1 day NH<sub>3</sub> | Yes | 6 days | 1 day |
| 1 day PBS | Yes | 1 day | - |
| 7 days PBS | Yes | 7 days | - |

### 2.4. X-Ray Microtomography (Micro-CT) analysis

This technique aimed to monitor the internal structure of collagen hydrogels. Gels were loaded with Micropaque contrast agent (Guerbet) before their observation and scanned using a high-resolution X-ray micro-CT system (Quantum FX Caliper, Life Sciences, Perkin Elmer, Waltham, MA, United States) hosted by the PIV Platform (UR2496, Montrouge, France). Standard acquisition settings were applied (voltage 90 kV, intensity 160 mA), and scans were performed with a field of view of 1 cm^2^. Micro-CT datasets were analyzed using the built-in multiplanar reconstruction tool, Osirix Lite (Pixmeo, Switzerland), to obtain time series of images and 3D reconstruction.

### 2.5. Second-harmonic generation (SHG)

Second-harmonic generation microscopy was used to observe anisotropy as a strong SHG signal is proportional to collagen quantity and alignment. SHG images were acquired with a Mai Tai multiphoton laser-equipped confocal microscope (Leica SP8). SHG signal was collected with a hybrid detector between 430 and 450 nm (excitation at 880 nm). A long-distance objective (25X water immersion) was used to acquire z-stacks with optimal settings for each sample. Image series were acquired with the Leica Application Suite X software.

### 2.6. Polarized Light Microscopy

Collagen anisotropy of 3D-printed hydrogels was also analyzed by polarized light microscopy (PLM). PLM was performed using a photonic Nixon microscope equipped with crossed-polarizers to observe collagen birefringence. Collagen fibrils with uniaxial alignment will increase or decrease the light intensity depending on the relative orientation of the crossed polarizer to the sample. A series of images were acquired with multiple sample orientations to demonstrate the light intensity variation. Maxima and minima were separated by 45°. These observations were correlated with those obtained by SHG.

### 2.7. Rheological measurements

Shear oscillatory measurements were performed on casted collagen hydrogels with different gelling parameters using an Anton Paar rheometer. An 8 mm parallel-plate geometry was fitted with a rough surface to avoid gel slipping. All measurements were performed at 37°C. Storage modulus, G’ and loss modulus, G” were recorded during a frequency sweep from 0.1 to 10 Hz with an imposed strain of 1%. This strain corresponds to non-destructive conditions (linear viscoelastic regime) as previously checked (data not shown). The geometry and a stabilized normal force (0.01 N) were optimized. Three samples of each matrix were tested.

### 2.8. Tensile tests

Mechanical tests were performed on printed gels with a tensile testing machine (Instron) equipped with a 10 N load cell and Bluehill software. Printed gels were obtained by printing 5 successive layers of 25 mm x 5 mm either in the longitudinal or transversal direction. All experiments were performed on hydrated samples (three specimens per condition) at 21°C at a constant strain rate of 0.06 s^-1^. The elastic (Young’s) modulus of collagen hydrogels was measured by a linear fit within the first 5% deformation.

### 2.9. Transmission Electron Microscopy (TEM)

For one hour, collagen hydrogels were fixed using 3.63% glutaraldehyde in cacodylate/saccharose buffer (0.05 M/0.3 M, pH=7.4). Samples were washed three times in cacodylate/saccharose buffer (0.05 M/0.3 M, pH=7.4). Samples were colored with nuclear red for 1 hour before progressive dehydration through increasing baths of ethanol and propylene oxide. Last, the hydrogels were embedded in Araldite. Thin Araldite transverse ultra-thin sections (70 nm) were performed using a Leica EM UC7 ultramicrotome and contrasted with 0.5% (w/v) uranyl acetate. Sections were then observed with a Cryo-microscope Tecnai spirit G2 electron microscope 214 operating at 120 kV. For each 3D-printed or cast hydrogel, photos were taken at a magnification of x15,000 with a 215 CCD Camera (Orius Gatan 832 digital) and analyzed.

### 2.10. Differential Scanning Calorimetry (DSC)

10-20 mg of collagen hydrogel was placed into an aluminum pan. The reference was an empty pan. Measurements were acquired with a NanoScan Differential Scanning Calorimetry. A temperature scanning from 20°C to 100°C with a 10°C.min^-1^ ramp was performed to detect the collagen fibril denaturation peak.

### 2.11. Fibroblast cell culture to observe collagen anisotropy

Normal Human Dermal Fibroblasts (NHDF) were used for their ability to sense the substrate topography. They are a living control for the anisotropy induced in the collagen constructs as they align along the longitudinal fibril axis. Briefly, they were cultured in a complete cell culture medium (Dulbecco’s Modified Eagle’s Medium (DMEM) supplemented with 10% fetal bovine serum, 100 U.mL^-1^ penicillin, 100 μg.mL^-1^ streptomycin, 0.25 μg.mL^-1^ Fungizone, and Glutamax) for 48 hours. Then, they were trypsinized (with 0.1% trypsin and 0.02% EDTA), and 5,000 cells were seeded on top of collagen hydrogels and cultivated for one day to observe their alignment. After fixation with paraformaldehyde, 4% overnight at 4°C, and staining with Phalloidin Alexa fluor 488 nm, hydrogels were analyzed with SHG coupled with fluorescent microscopy. Fibroblasts orientation along the collagen filaments was analyzed using the Orientation J plug-in (Fiji).

### 2.12. Pore generation

Large channels were added to the scaffolds using needles. First, the hydrogels were printed into a PBS 5X bath or in the air (for the negative control) inside dedicated 3D-printed plastic molds with holes (10 × 5 × 1.5 mm). After 30 min of gelling inside PBS 5X, 2 needles (external diameter 600 µm/23G) were placed into the collagen scaffold through the dedicated holes of the mold. The fibrillogenesis and gelling processes were then continued. After complete gelling and a PBS 1X rinsing, needles were removed to reveal straight, well-defined channels that cross the whole hydrogel.

### 2.13. C2C12 culture inside large channels

C2C12 were expanded in proliferation medium (DMEM high glucose, 20% fetal bovine serum, 100 U.mL^-1^ penicillin, 100 μg.mL^-1^ streptomycin) until 50% confluency. They were trypsinized and amplified to avoid confluency.

To seed C2C12 into printed collagen hydrogels (1 day PBS/1 day NH_3_), cells were resuspended in Matrigel at 3.10^7^ cells.ml^-1^ (Sigma, Bioreagent) on ice. 3 µl of cell suspension were injected with a P10 pipet in each channel created by needles (600 µm). After 24h, two conditions were used: a first group was left in proliferation medium for 4 or 7 days in culture, whereas the other group was cultivated during the same period with differentiation medium (DMEM high glucose, 2% horse serum, 100 U.mL-1 penicillin, 100 μg.mL-1 streptomycin). The medium was changed every day. Constructs were then fixed in paraformaldehyde 4% in PBS 1X overnight at 4°C and sectioned into 250 µm slices with a vibratome. All experiments were carried out in triplicates.

### 2.14. Fluorescence and Second-Harmonic Generation microscopy

For constructs colonized by C2C12, the fluorescent labeling of nuclei (TOPRO-3, Thermofisher) and actin filaments (Alexa Fluor 488 phalloidin, Thermofisher) were performed on 250 µm sections containing the channels. Additional labeling of C2C12 with MF-20 hybridoma mouse IgG2B primary antibody and Alexa Fluor 546 goat anti-mouse IgG2B secondary antibody (Invitrogen) was used to evaluate the cell differentiation into myotubes. Observations were conducted with a Leica SP5 upright confocal, multiphoton laser scanning microscopy, which enabled the simultaneous acquisition of fluorescence and second-harmonic generation signals.

### 2.15. Statistical analysis

All experiments were carried out at least twice, and the results were expressed as the mean values ± standard deviation (SD). In all cases, data are means ± SD. The differences were analyzed using Mann-Whitney tests (when n>4), *p* < 0.05 was considered significant.

## 3. Results

### 3.1.3D printing of porous dense collagen hydrogels

The home-built 3D printer enabled the printing of 30 mg.mL^-1^ collagen solutions through a 23G flat bottom needle, creating a constant flow of collagen solution. The printer head extruded the collagen through the needle, and collagen molecules aligned along the axis of the needle during the process (Figure 1-A). However, this printing is only possible with suitable parameters such as printing speed, printing condition (air or PBS 5X), or height between two successive layers. First, the printing speed was varied from 1 to 10 mm.s^-1^, but 2 mm.s^-1^ was chosen to obtain a homogeneous thread without collagen excess. Then, the filament was either printed in a petri dish filled with PBS 5X (for all conditions except 1 day NH_3_) or in the air (for 1 day NH_3_ gelling). The PBS 5X bath gelled quasi-instantaneously the collagen solution and allowed an excellent shape fidelity and reproducibility of the printing process (Figure 1-B and C). Finally, to tune the space between two layers, the same layer was repeated at different z-positions. Gelling kinetics in PBS 5X was founded to be the best compromise for a printing bath: sufficiently fast to preserve the round shape of filaments (and create intrinsic porosity between filaments) and slow enough to ensure the scaffold function by melting adjacent filaments together (Figure 2-A, B and C).

**Figure 1.**
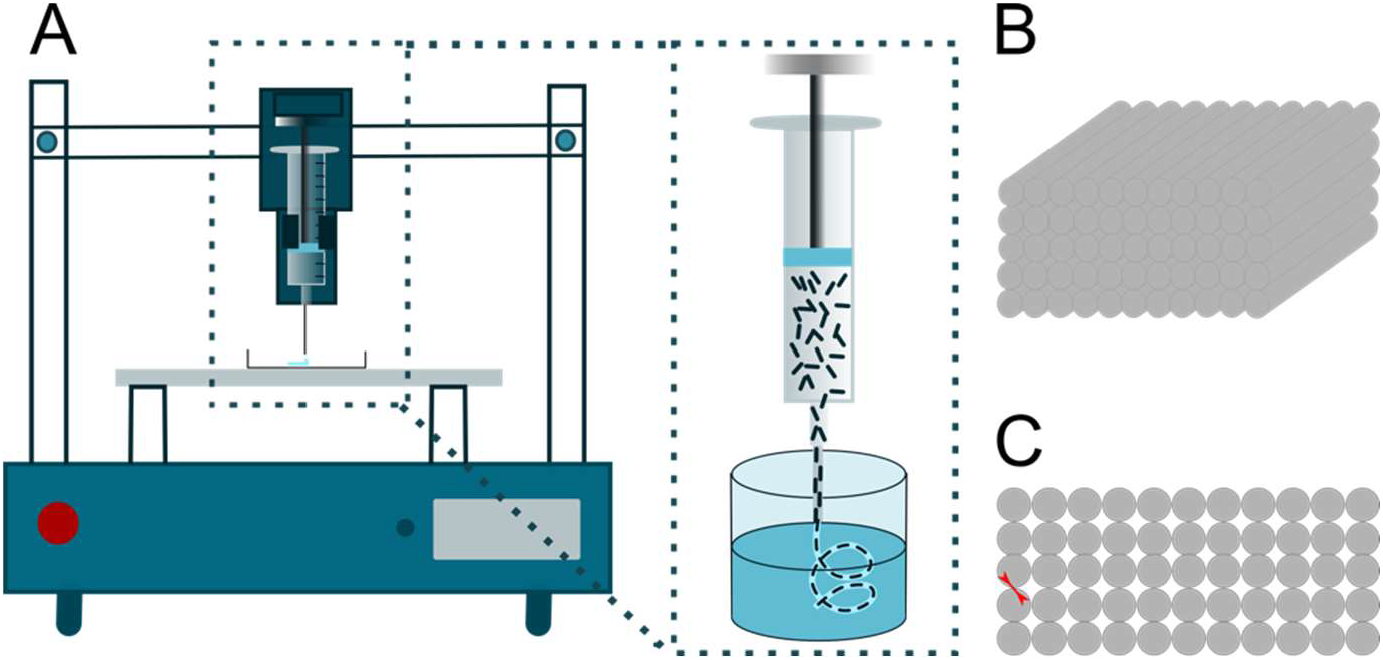
(A) 3D printing set up. The dense collagen solution was extruded through a 23G flat bottom needle. Collagen molecules aligned along the axis of the needle and the PBS 5X bath froze their anisotropic organization thanks to the rise in pH and the osmotic pressure. (B) 3D view and (C) Lateral view of the printed gel (gel dimension 1 cm x 1 cm x 2 mm. The black arrow indicates the interlayer space, and the red arrow indicates the diameter of the pores.

**Figure 2.**
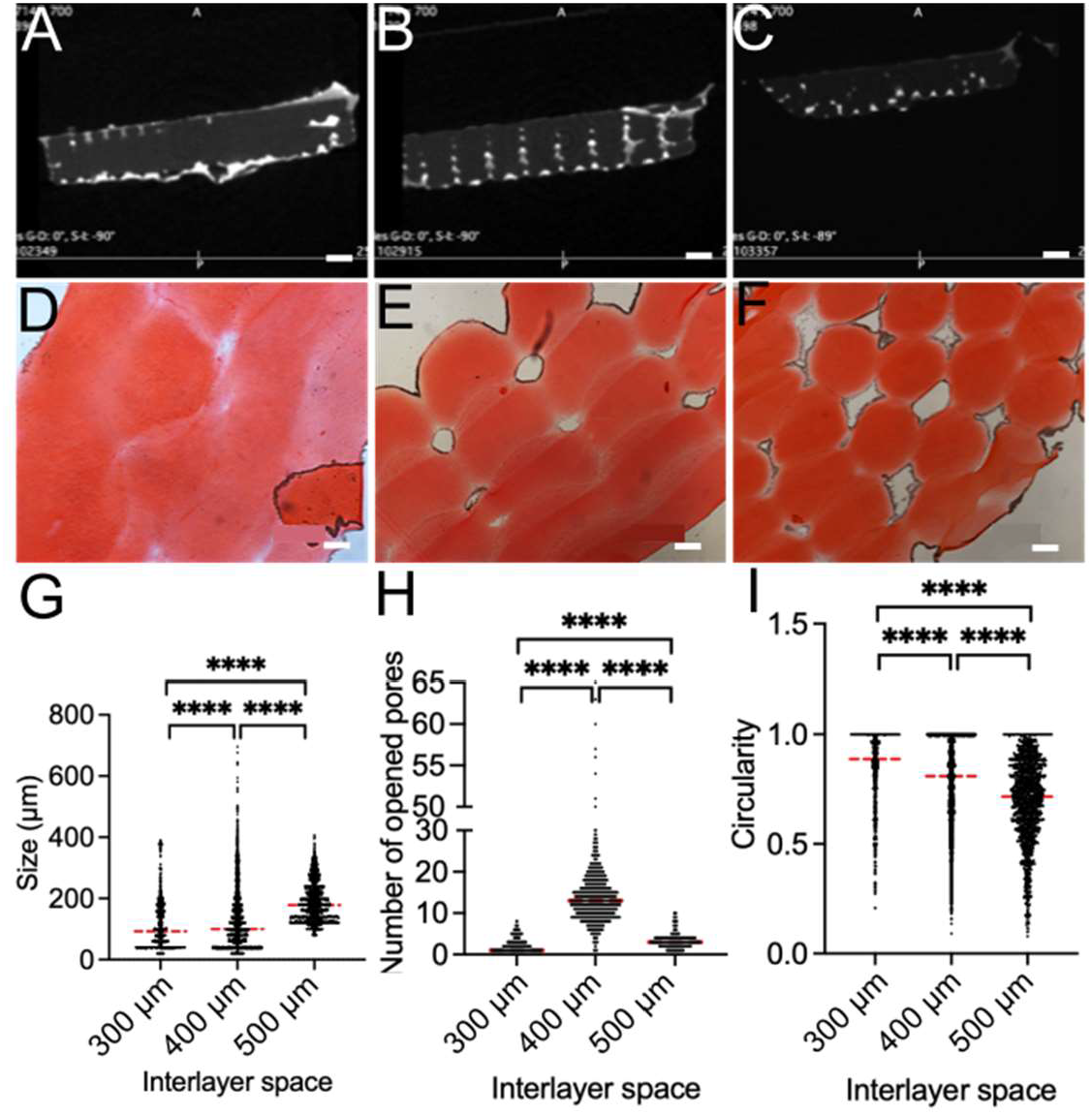
Intrinsic porosity of 3D-printed collagen hydrogels, a stack of layers made of 330 µm thick filaments. The top layer is cohesive with the previous ones, generating an inherent porosity. (A-C) 3D-printed hydrogels observed by micro-CT (scale bar 1 mm) Interlayer space: (A) 300 µm, (B) 400 µm, and (C) 500 µm. (D-F) Sirius red staining on histological sections from printed collagen hydrogels (scale bar 100 µm). Interlayer space: (D) 300 µm, (E) 400 µm and (F) 500 µm. (G) Image analysis of pore size distribution depending on the interlayer space (Feret min diameter of pores). (H) Image analysis of pores number depending on the interlayer space. (I) Image analysis of pores circularity (Red lines indicate the median,****: *p* < 0.0001, Mann-Whitney test).

The generated porosity depends on the space between each printed layer and can reach a diameter of 150 µm for the 500 µm interspace (Figure 2-G). The number of pores also depended on the interlayer space and was optimal for 400 µm (Figure 2-H) as many pores collapsed for 300 µm and 500 µm. The hydrogel cohesiveness decreased with a 500 µm interlayer space, and the well-organized stack was lost. This explains the decrease in pore circularity and the low number of open pores (Figure 2-H and I). From these different tests, the best combination between cohesiveness and porosity was found for an interspace of 400 µm between each layer, generating well-dispersed 100 µm intrinsic pores that cross the hydrogel from one side to the other (Figure S1 and S2). Following the printing process in the initial PBS 5X bath, the fibrillogenesis process was continued in PBS 5X or under ammonia vapors according to the different tested conditions (see Table 1). Once gelled, extensive washes in PBS 1X restored a neutral pH compatible with cell culture, and constructs were stored in PBS 1X at 4°C until use.

### 3.2. Evaluation of collagen anisotropy

Different techniques were used to assess collagen anisotropy from a macroscopic aspect (second-harmonic generation (SHG) and polarized-light microscopies (PLM)) to a microscopic aspect with cell alignment along collagen fibers. First, printed hydrogels exhibited different SHG signal intensities according to the gelling method for the same quantity of collagen (see Table 1). Printed collagen constructs gelled with ammonia vapors alone (1 day NH_3_) or combined with PBS 5X for 30 min (30 min PBS/1 day 1 day NH_3_) exhibited a weak SHG intensity compared to other constructs (Figure 3 – row A), showing the disorganization of collagen fibrils. The analysis under crossed-polarized light microscopy confirmed the results obtained by SHG (Figure 3 – row B). Collagen birefringence was visible for all the conditions gelled in PBS 5X for at least one day.

**Figure 3.**
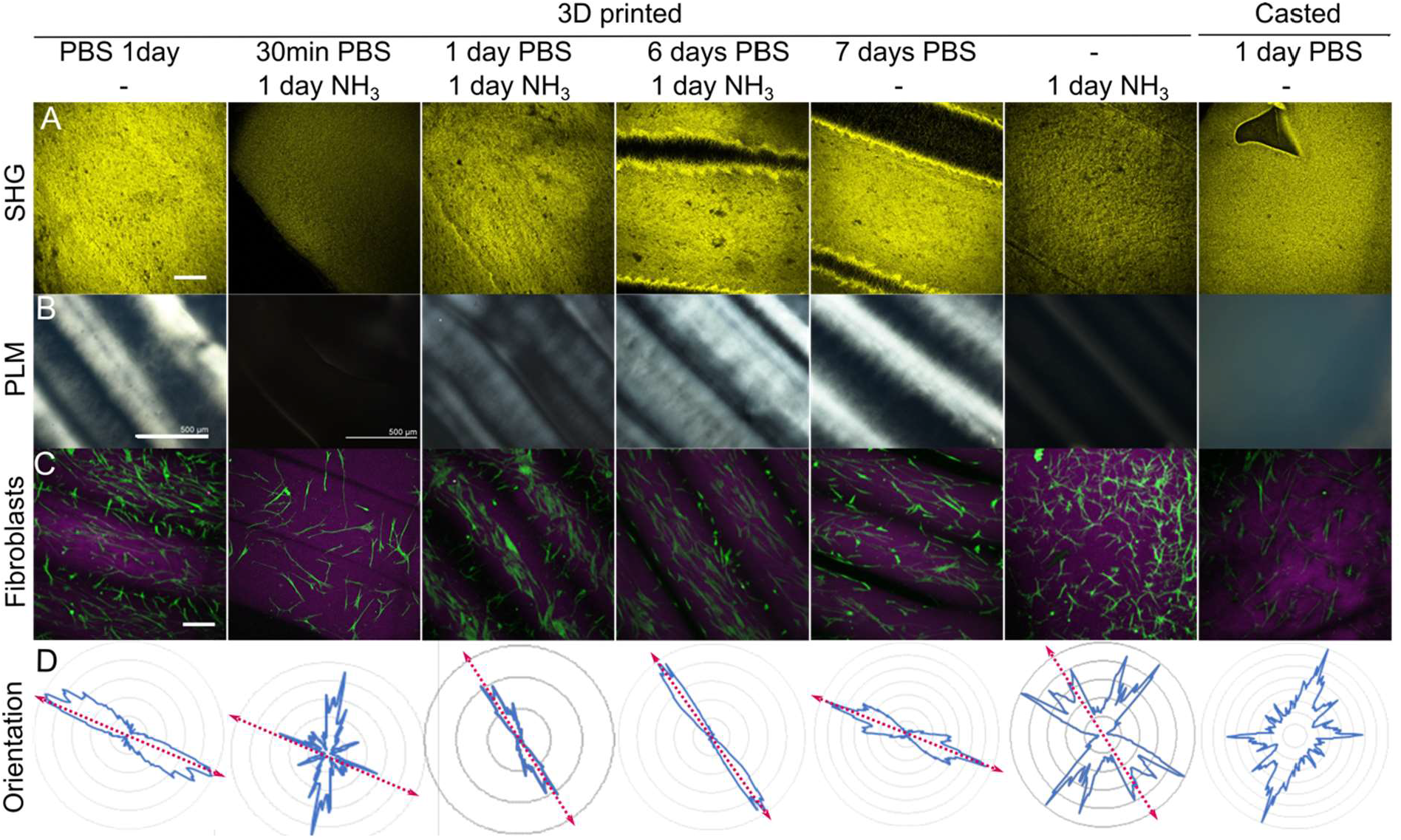
Anisotropy observation and quantification within 3D-printed dense collagen hydrogels. (A) Second-harmonic Microscopy (SHG) Imaging (scale bar 100 µm). (B) Polarized Light microscopy (PLM) (scale bar 500 µm). (C) Fibroblasts alignment on hydrogels. Green: Fibroblasts labeled with Alexa Fluor 488 Phalloidin, Purple: SHG signal from collagen fibrils. (scale bar 250 µm). (D) Quantification of cell alignment by Orientation J treatment. Percentage of cells depending on the angle they made with the filament orientation (0°). The pink arrow indicates the filament orientation.

Conversely, casted hydrogels did not exhibit collagen birefringence regardless of the gelling process (Figure 3 -last column and Figure S3). Last, the presence of anisotropy was investigated by the culture of fibroblasts at the hydrogel surface. After 24 hours in culture, fibroblasts aligned along the axis of the filaments of 3D-printed hydrogels (Figure 3 – row C). The graphs obtained with the Orientation J plug-in (Fiji) demonstrated that all printed conditions, except 1 day NH_3_ and 30 min PBS/1 day NH_3_, were suitable for fibroblast alignment (Figure 3 – row D). In contrast, fibroblasts were randomly oriented when seeded onto casted hydrogels (Figure 3 - last column and Figure S3). Hence, a minimum of 24 hours in PBS 5X was required to maintain the anisotropy created by the printing process.

### 3.3. Impact of 3D printing on collagen mechanical properties

To monitor the impact of the gelling process on mechanical properties, rheology was first performed on the different casted hydrogels. 1 day of NH_3_ gelling led to the formation of hydrogels with a high storage modulus, G’ around 2.5 kPa. The combination of gelling methods 1 day PBS/ 1 day NH_3_ generated hydrogels with a similar storage modulus. Hydrogels only gelled with PBS 5X (1 day PBS, 7 days PBS) exhibited a lower stiffness with G’ around 1 kPa. The other conditions combining PBS 5X and NH_3_, *i*.*e*., 30 min PBS/1 day NH_3_ and 6 days PBS/1 day NH_3,_ did not permit the generation of hydrogels with high mechanical properties (Figure 4-A). Rheology was suitable for casted isotropic gels but was irrelevant to printed gels due to their anisotropy. Thus, tensile tests were performed on printed hydrogels (along fiber direction and perpendicular to fiber direction) to observe the impact of printing and anisotropy on the mechanical response. For 1 day PBS constructs, the stretching axis dramatically changed the mechanical properties, thereby evidencing anisotropy presence (Figure 4-B). A 1 day PBS/1 day NH_3_ gelling reproduced the same mechanical behavior, *i*.*e*., different curves between the traction along the fibrils and perpendicularly (Figure 4-C). However, the difference was lower. Last, 1 day NH_3_ constructs, the curves completely overlaid, showing no significant differences between the two orientations of stretching. This evidences the absence of anisotropy (Figure 4-D). The Young’s modulus obtained by a linear fit on the first 5% deformation did not reveal significant differences between 1 day PBS and 1 day PBS/1 day NH_3_ constructs (Figure S4).

**Figure 4.**
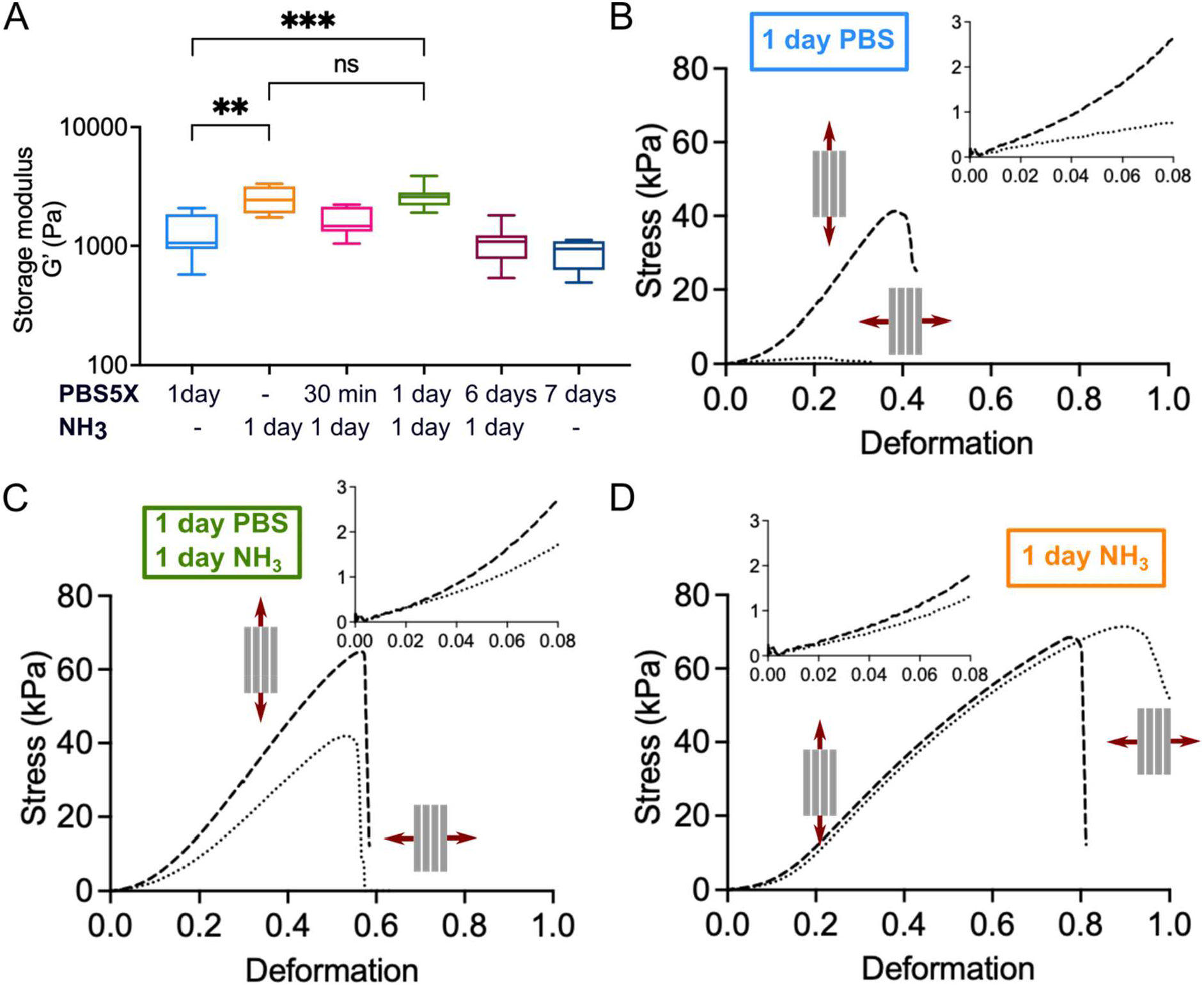
(A) Storage Modulus (G’) of dense collagen hydrogels assessed by rheological measurements. (***: *p* < 0.001, **: *p* < 0.01, ns: *p* > 0,05 Mann-Whitney test). (B-D) Tensile test curves of printed collagen construct stretched along the fibrils (non-continuous line) and perpendicularly (dotted line). Experiments were performed for (B) 1 day PBS, (C) 1 day PBS/1 day NH_3_, and (D) 1 day NH_3_.

### 3.4. Fibrillar structure of collagen hydrogels

Collagen fibrillar structure was analyzed by Transmission Electron Microscopy (TEM) to observe the impact of the printing process and gelling conditions on fibrillogenesis. Using ImageJ, a surface density of fibrils was measured on TEM images to compare fibrillogenesis in printed and casted hydrogels. This analysis showed that the fibril density measured in printed gels formed with PBS (1 day PBS or 7 days PBS) or with mixed conditions (30 min PBS/1 day NH_3_, 1 day PBS/1 day NH_3_, 6 days PBS/1 day NH_3_) was higher than that in casted hydrogels (Figure 5-A, B and C). This increase in fibril density was not visible for 1 day of NH_3_ gelling condition. The differential calorimetry experiment did not demonstrate any modifications in the temperature of the denaturation peak of collagen, whatever the gelling condition. Thus, all conditions formed mature collagen fibrils (Figure 5-D).

**Figure 5.**
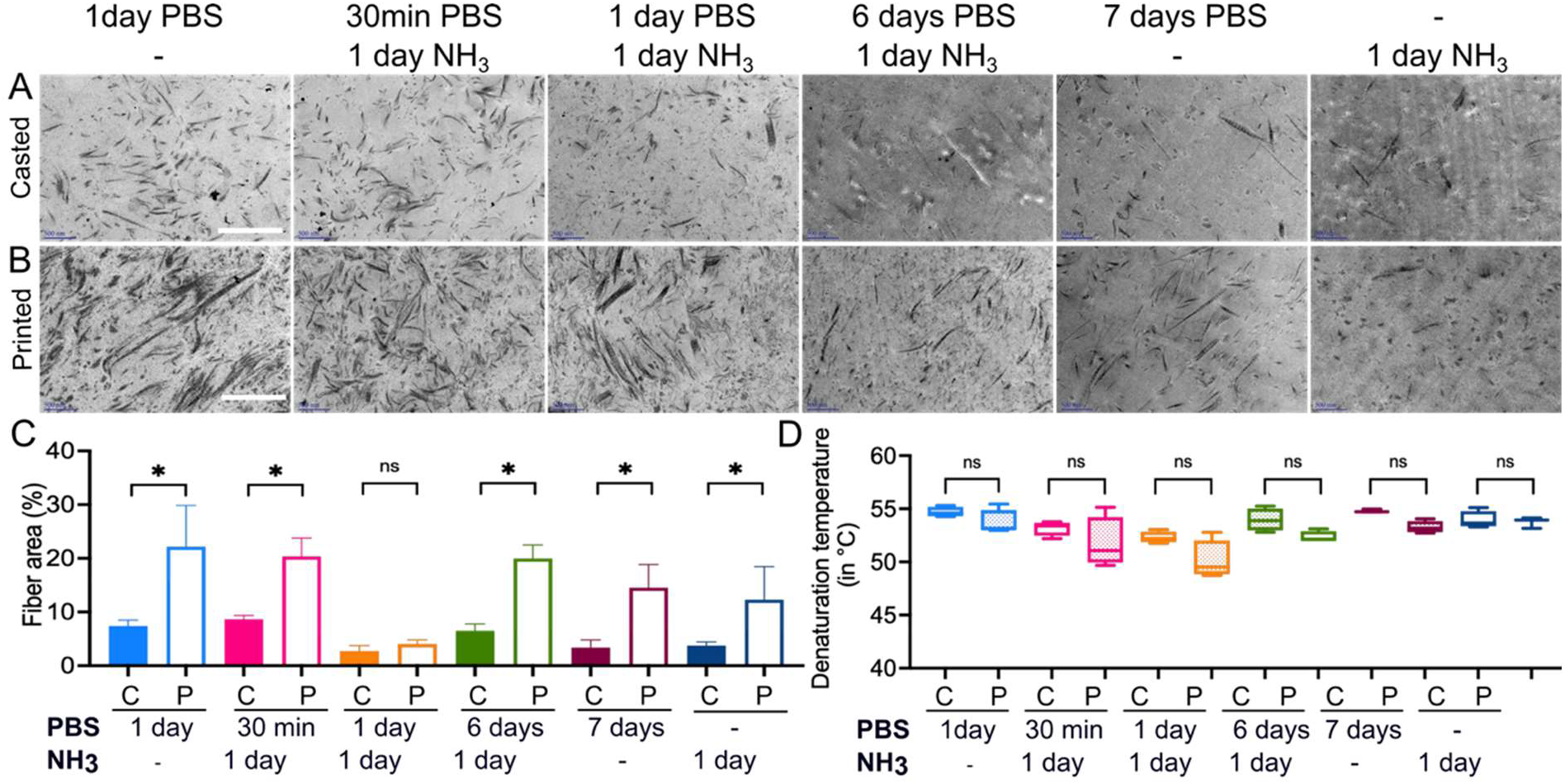
Collagen fibrillogenesis according to the gelling conditions observed by Transmission Electron Microscopy. (A) images of casted hydrogels (scale bar 500 nm). (B) 3D-printed hydrogels (scale bar 500 nm). (C) Analysis of the area covered by fibrils in casted (C) and printed (P) gels. D: Denaturation temperature of collagen fibrils within casted (C) or 3D-printed (P) hydrogels regarding the gelling process.

### 3.5. Addition of large channels within the collagen scaffold for cell colonization

*In vivo*, muscle bundles are more significant than the 100 µm created by the stacking of filaments. Hence, an additional porosity is required for cell cultivation. The best combination between anisotropy and mechanical properties was 1 day PBS/1 day NH_3_ and was selected for the rest of the study. To have reproducible hydrogels, a dedicated plastic mold was created to place the needles in the same position (Figure 6-A). Collagen constructs were directly printed inside this mold. After a 30 min period post-printing in PBS 5X, forming gels were perforated with 600 µm needles using the dedicated holes. Then, the gelling was extended with 1 day PBS/1 day NH_3_ to increase mechanical properties. After complete gelling, micro Computed Tomography revealed 2 well-defined straight channels inside the printed hydrogels (Figure 6-B). The smaller intrinsic pores obtained by the printing process were also visible on the scan. Collagen birefringence was still visible. The SHG signal was also not altered (Figure 6-C et D), indicating that the addition of needles post-printing did not disturb the anisotropy generated during the first short gelling period in PBS 5X (Figure S5).

**Figure 6.**
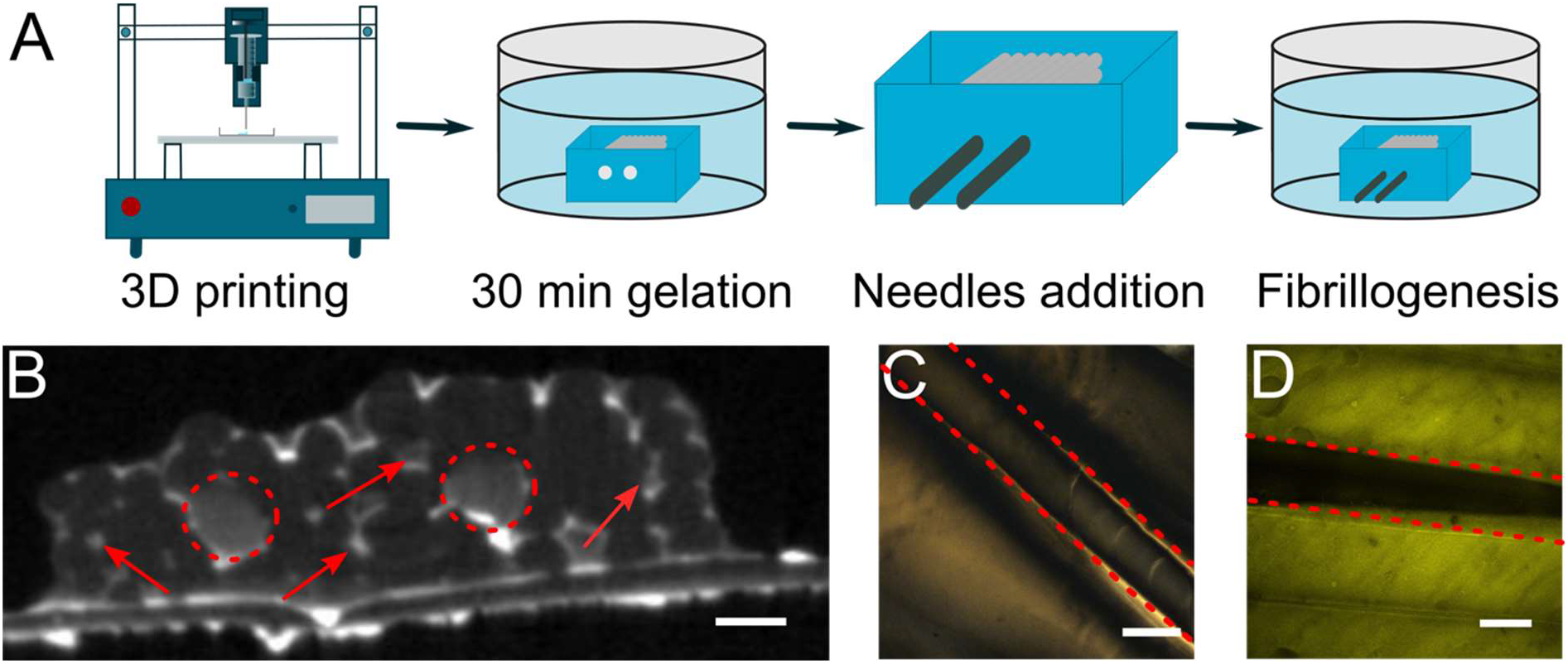
Generation of large channels within printed hydrogels without alteration of collagen anisotropy. (A) Set up: needles molding during collagen gelation. (B) Micro Computed Tomography of a 3D-printed gel with 2 needles. Two networks of pores were observed: one around 100 µm thanks to printing (intrinsic porosity – red arrows) and the other at 600 µm (needles – red circles) (scale bar 500 µm). (C) Cross-polarized light microscopy of sections of printed hydrogel (1 day PBS/1 day NH_3_) with the presence of needles. (scale bar 500 µm). (D) Second-harmonic generation imaging of sections of printed gel with needles (scale bar 250 µm).

### 3.6. Murine skeletal myoblast colonization and differentiation inside large channels

Myoblasts were seeded inside the large channels created within the perimysium-like scaffold to model muscle fibers. Muscle fibers are composed of myoblasts surrounded by loose connective tissue made of collagen IV and laminin. The use of Matrigel® reproduced the latter. C2C12 murine skeletal myoblasts were seeded and cultured for 4 or 7 days. Regardless of the medium used, proliferation, or differentiation, myoblasts aligned along the axis of the pores and filled the channel. When cultured in a differentiation medium, some parallel, multinucleated densely-packed myotubes expressing heavy myosin chains (MF20) could be observed on day 4, reproducing a muscular bundle. Fused myoblasts forming myotubes were easily visible with phalloidin staining thanks to their enlarged shape and numerous nuclei (Figure 7). Seven days in the differentiation medium did not show significant changes in differentiation (Figure S6).

**Figure 7.**
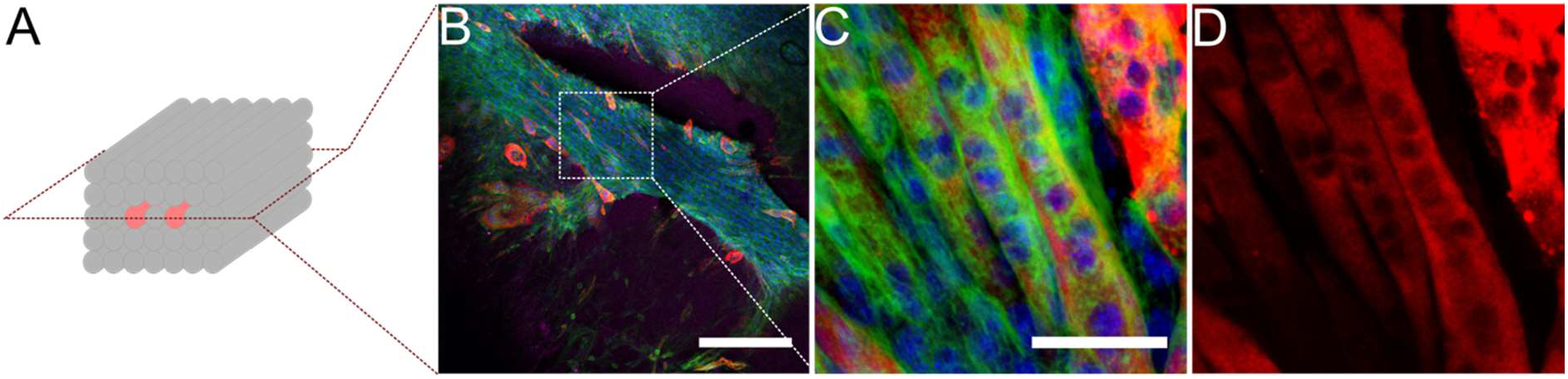
C2C12 colonization and differentiation into myotubes within the large pores generated by needles (600 µm). (A) Slicing axis for immunostainings. (B) Longitudinal view of pores (scale bar 100 µm). Green: actin, blue: nucleus, red: MF20. (C) Zoom on myotubes alignment and (D) myosin staining (scale bar 50 µm)

## 4. Discussion

This study aimed to design a novel *in vitro* 3D model of skeletal muscle combining a physiologically relevant artificial matrix with muscle bundles. A scaffold in the form of a dense collagen hydrogel was synthesized to mimic the perimysium. This synthetic matrix had to mimic the four main characteristics of the native one, *i*.*e*., its biochemical composition, stiffness, porosity, and anisotropy. Then, this scaffold was cellularized with C2C12 myoblasts embedded inside Matrigel® to reproduce the muscle bundles (myotubes + endomysium). After *in situ* differentiation, cells assembled into mature multinucleated myotubes.

First, the perimysium is mainly composed of collagen I and III, proteoglycans, and glycosaminoglycans [33]. The fibril-forming collagen I is a building block of great interest in fabricating the scaffold. This protein is known to strengthen connective tissues and is the natural support for cell adhesion. Unfortunately, collagen-based hydrogels are currently manufactured from low concentrated solutions (<4 mg.mL^-1^), possess poor mechanical properties, and shrink under cellular contractile activities [11,34]. These drawbacks can be circumvented without any crosslinking by increasing the collagen concentration. This study used evaporation to obtain a printable collagen solution at 30 mg.mL^-1^ [35], mimicking the physiological concentration observed in muscle ECM [36]. Despite the high viscosity of concentrated solutions, their printing at 2 mm.s^-1^ was accurate thanks to the shear-thinning behavior of collagen I [16,28]. It is consistent with the study of Rhee and co-workers showing that a collagen concentration superior to 17 mg.mL^-1^ is required to obtain printing accuracy and shape fidelity [20]. Unfortunately, the round shape of these extruded filaments rapidly disappeared in contact with air and the other filaments. Based on Picaut and co-workers’ study, a printing bath made of PBS 5X enabled good shape fidelity and rapid gelling [16]. The diameter of extruded threads remained stable (330 µm) without swelling or shrinkage, thanks to the appropriate ionic strength. Unlike the printing bath described by Lee and co-workers, based on gelatin particles, using PBS 5X does not require a heating step detrimental to collagen. It allows the easy printing of successive layers [19,37,38]. In addition, thanks to the interlayer space used, a cohesive and well organized structure was generated.

The muscular perimysium topography relies on collagen anisotropy. In these tissues, fibrils are highly oriented in the same direction to promote mechanotransduction inside the skeletal muscle [33]. In our study, the extrusion process creates an intrinsic anisotropy by aligning collagen molecules due to the shearing stress the needle applied to the collagen. This alignment is “frozen” by the PBS 5X bath, which rapidly diffuses (in a minute) within the whole volume of the thin collagen filaments and neutralizes acetic acid. Consequently, intrinsic anisotropy is maintained in dense hydrogels thanks to collagen fibrils forming in place of collagen triple helices.. Picaut demonstrated that a concentration superior to 30 mg.ml^-1^ within a printing bath containing PBS 5X permitted to obtain isolated anisotropic threads. However, 3D cohesive and complex constructs made of anisotropic and dense filaments of pure collagen I with a controlled concentration had not been described yet. Collagen anisotropy of 3D cohesive constructs is often obtained after printing composite materials consisting of decellularized cornea matrix, collagen/Pluronic 127, or a collagen/hyaluronic acid to ensure a mechanical strength [39–41]. The major drawbacks of these systems are the absence of biomimetic collagen fibrils, the significant volume loss after polymer removal, or the strong collagen interaction with the other polymers, thereby preventing an adequate interaction scaffold/cells. In this study, pure dense collagen was very close to the physiological ECM structure and promoted muscle bundles-perimysium interactions.

After printing, an extended gelling time was required to obtain a complete collagen fibrillogenesis and suitable mechanical properties. During collagen gelling, two phenomena compete: collagen lateral self-assembling into fibrils (fibrillogenesis) and forming physical crosslinking nodes between collagen molecules to get a hydrogel [32]. Fibrillogenesis is strongly impacted by collagen concentration, ionic strength, and pH [42]. In this work, two methods were used for the extended gelling: (1) gelling with PBS 5X (pH 7.4, high ionic strength) and (2) NH_3_ vapors (pH 11, low ionic strength). PBS gelling starts immediately, but fibrils length and thickness evolve across time, whereas NH_3_ vapors trigger a rapid gelling after a lag phase (time for NH_3_ to dissolve and diffuse into the collagen solution). The high ionic strength promotes fibril growth, which explains the larger diameter of fibrils observed in PBS 5X compared to NH_3_. [32]. However, fibrillogenesis occurred efficiently in all conditions as a denaturation peak at 55°C was always observed [43].

The pH highly impacts collagen hydrogel mechanical properties during the gelling method [42]. At pH 10-11, collagen molecules are highly charged, and electrostatic interactions are favored to form physical crosslinking junctions, but hydrophobic interactions required for collagen fibrillogenesis are inhibited. This explains the high mechanical properties and the low fibril diameter observed in hydrogels only gelled with NH_3_ [32,35]. On the contrary, PBS gelling favors fibril growth but not physical crosslinking, leading to lower mechanical properties.

The traction tests performed along with the fibril orientation and perpendicularly to this orientation revealed different mechanical behaviors for the 1 day PBS and 1 day PBS/1 day NH_3_ constructs. Unlike hydrogels formed with NH_3_, the tensile test curves proved the anisotropy of these constructs as the tensile curves differ according to the stretching axis, impacting Young’s modulus.

The condition leading to a biomimetic muscle ECM mimicking perimysium was 1 day PBS/1 day NH_3_ because it combines the fibril formation and the appropriate mechanical properties. When 1 day PBS/1 day NH_3_ printed constructs were stretched in the fibril direction, Young’s modulus of hydrogels gelled increased up to 25.1 ± 9.6 kPa but slightly decreased down to 16.01 ± 7.6kPa when extended perpendicularly (Figure S4). All these values remain in the range of stiffness (8 – 25 kPa), promoting myogenic differentiation [44]. In addition, the mechanical behavior is elastic when a 10% deformation is applied, as seen on traction curves. For the first time, by combining two methods of collagen gelling (PBS 5X and NH_3_), we obtained anisotropic collagen constructs made of large biomimetic fibrils and exhibiting the mechanical behavior of the muscle ECM [45].

An anisotropic dense collagen hydrogel is not sufficient to mimic muscle extracellular matrix. Porosity is significant since it represents 70% of the global decellularized volume [33]. *In vivo*, muscle cells fill these pores to form muscle bundles from mm to cm long [46]. These muscle fibers are surrounded by the perimysium, in which a dense capillary network brings oxygen and nutrients to the cells [47]. This capillary density is required to meet the oxygen and nutrient demands as cells must not exceed 200 µm to survive [10]. *In vitro*, diffusion is insufficient to ensure nutrients and oxygen diffusion into a compact 3D dense collagen construct. The 3D printing process of our study created an adequate intrinsic porosity between the different collagen layers. Thanks to the high shape fidelity, extruded threads remain round and will gently be superposed on top of each other. During the initial gelling, the filaments partially fused with their neighbors to allow the construct cohesiveness. From a side view, the construct resembles a “stack of wood” with interstitial space between every filament. This one-step process generates a porosity of 100 µm in diameter cylindrical pores crossing the anisotropic construct and is suitable for nutrients and oxygen diffusion.

Meanwhile, in this condition, the cohesiveness of the construct was preserved. This porosity generation does not require additional steps such as heating compared to sacrificial ink techniques or porogen leaching. Unlike the bidirectional methods currently used in 3D printing (angle between one layer and the upper one 90°) to improve the construct cohesiveness, the straight channels generated by our procedure allowed for a rapid and effective liquid diffusion. [28].

This intrinsic porosity is adequate for nutrient diffusion but is too tiny for muscle cell colonization and organization. A muscle bundle comprises 20 to 60 muscle cells (20-100 µm in diameter) corresponding to a final diameter of 400 to 6000 µm [1]. A 600 µm diameter corresponds to the physiological range of a muscle bundle. Generating very large channels by 3D printing is difficult as channels tend to collapse. For this reason, we used a molding method. Two straight channels were easily created using 600 µm needles within dense collagen hydrogels without disturbing the collagen alignment after removal. Hence, Anisotropy appeared at two different scales: anisotropic channels (high length/width ratio), forcing myoblasts to align [31,48], and inherent anisotropy within channel walls (aligned fibrils of collagen due to the printing process).

C2C12 myoblasts were seeded into the large pores of printed hydrogels using the 1day PBS/1 day NH_3_ gelling condition to observe myotubes formation in a 3D biomimetic environment. Matrigel® mimicked the endomysium because of its similar composition, mostly collagen IV and laminins [33]. After 4 days of differentiation, large, densely-packed, and aligned multinucleated myotubes expressing myosin heavy chain were observed within the whole volume of channels. Hence, the scaffold was suitable to sustain the development of fused myoblasts into myotubes in 3D after 4 days. Previous works investigated the C2C12 behavior on different porous collagen structures [26,31]. Unfortunately, the 3D organization of cells in these materials was questionable because cells only adhered to the pore surface and did not fill up the entire volume to acquire an organotypic organization.

The model of skeletal muscle presented here is based on the development of a biomimetic ECM. Despite interesting cell contractility results, 2D muscle models do not study the cell-ECM interactions and lack 3D organotypic structure. 3D models seemed to bring a new approach, but the extracellular matrix role was often neglected. The most widespread 3D *in vitro* model was developed to study contractility. A low-concentrated collagen solution encapsulates cells in a 3D shape before creating a microtissue between PDMS pillars [7,8]. However, cells shrank the hydrogel and died because of a lack of nutrients and oxygen diffusion [49]. In addition, ECM physical properties are poorly tunable and controllable due to shrinkage. Another model used synthetic polymers to create large channels for myoblasts seeding [48,50]. This scaffold had high mechanical properties but failed to reproduce cell adhesion cues. Other approaches are based on the 3D printing of a cell-laden ink to create cellularized threads mimicking muscular bundles [21,22], but collagen is rarely used. Kim and co-workers developed bioprinted threads of dense collagen colonized by C2C12, but the diameter of their muscle bundle is at least twice lower compared to physiological bundles [17]. The poor nutrient diffusion inside dense collagen justifies this choice.

In our model, the biochemical cues are restored thanks to collagen I and Matrigel®. Our collagen constructs’ stiffness, porosity, and anisotropy imitate the muscle cell microenvironment.

## 5. Conclusions

In this study, we have designed a biomimetic hydrogel possessing the physical properties of the muscle ECM and colonized it with C2C12 myoblasts to form muscle bundles. The unidirectional 3D printing of dense collagen solutions within an appropriate gelling bath is an adequate strategy to mimic perimysium. This single-step technique produces dense anisotropic hydrogels possessing interesting mechanical properties and an intrinsic porosity suitable for nutrient and O_2_ diffusion. In addition, larger channels dedicated to cell colonization and muscle bundle organization can be easily set by molding without disturbing the hydrogel topography and mechanical properties. Last, this model allows the muscle cell alignment within channels filled with Matrigel® and their differentiation into myotubes, thereby reproducing the structure of a skeletal muscle bundle. Compared to 2D models, this 3D novel model brings more complexity despite its simple fabrication. It will permit to study of the impact of the muscle ECM properties on cell phenotype and contraction.

## Supporting information

Supplementary Data - Article Marie Camman

## Declaration of competing interests

The authors declare no conflict of interests relevant to this work.

## Notes

### Competing Interest Statement

The authors have declared no competing interest.

