## Supplementary Data - Article Marie Camman for "Anisotropic dense collagen hydrogels possessing two ranges of porosity to create the adequate microenvironment for muscle bundles: a step towards skeletal muscle modeling"

Postal address: Campus Pierre et Marie Curie, 4 place Jussieu, 75252 Paris Cedex 05, France

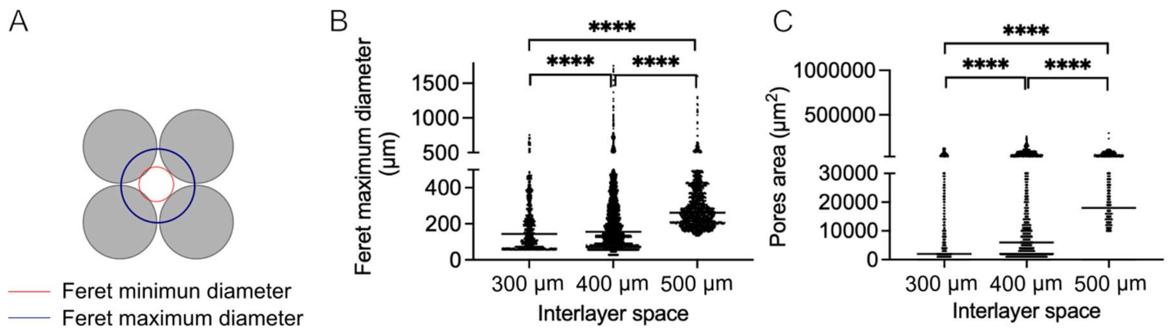


**Figure S1.** A- Determination of Feret diameter to evaluate pore size. B- Feret maximum diameter depending on the interlayer space. C- Pores area depending on the interlayer space.

| | 0.3 mm | 0.4 mm | 0.5 mm |
|---|---|---|---|
| Pores mean area (μm2) | 19440 ± 26210 | 23620 ± 34290 | 35190 ± 24770 |
| Feret min (μm) | 114,6 ± 84,87 | 127,5 ± 93,93 | 187,1 ± 64,56 |
| Feret max (μm) | 186,7 ± 137,2 | 205,5 ± 162,1 | 294,8 ± 139 |
| Circularity | 0,8339 ± 0,1861 | 0,7797 ± 0,1976 | 0,6923 ± 0,1972 |
| Number of pores | 1,719 ± 1,795 | 14,11 ± 7,034 | 3,203 ± 1,929 |

**Figure S2.** Table of numerical results after image analysis.

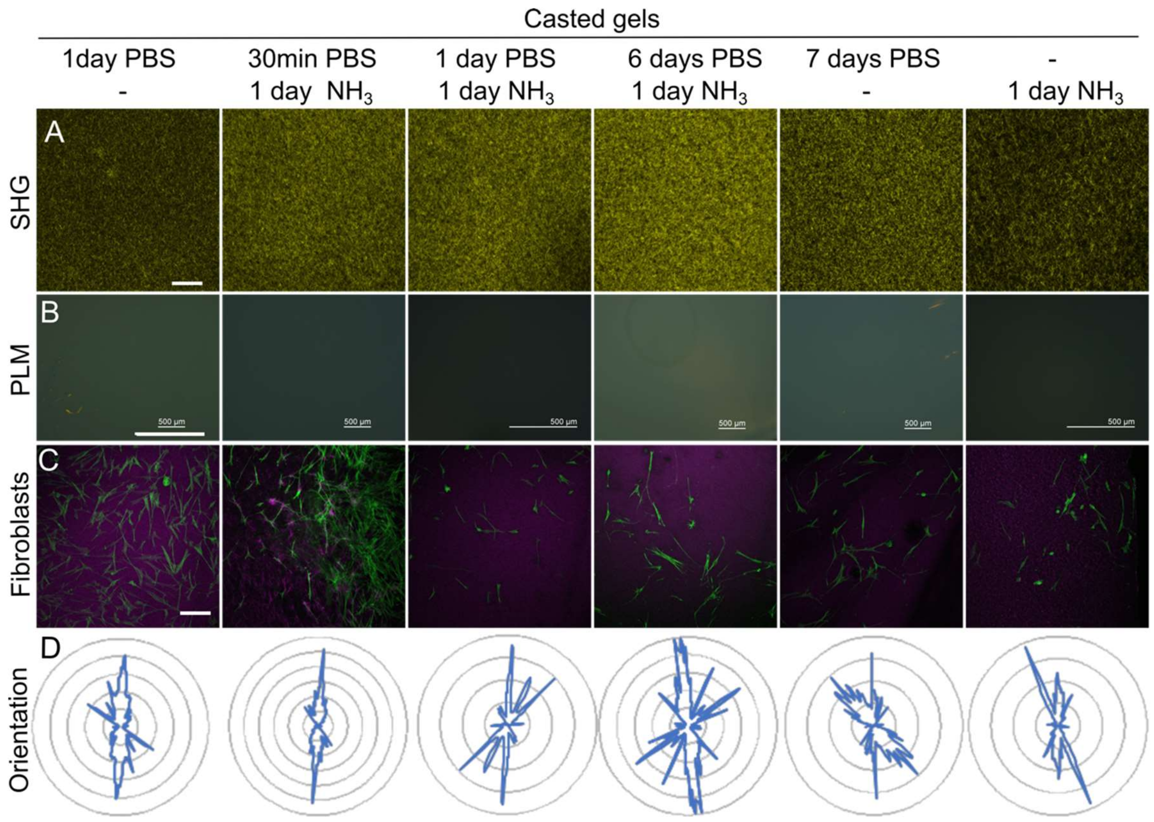


**Figure S3.** Anisotropy observation and quantification within casted dense collagen hydrogels. (A) Second-harmonic Microscopy Imaging (scale bar 100 µm). (B) Polarized Light microscopy (scale bar 500 µm). (C) Fibroblasts alignment on hydrogels. Green: Fibroblasts labeled with Alexa Fluor 488 Phalloidin, Purple: SHG signal from collagen fibrils. (scale bar 250 µm). (D) Quantification of cell alignment by Orientation J treatment. Percentage of cells depending on the angle they made with the filament orientation (0°).

| Young's modulus (kPa) | 1 day PBS | 30 min PBS<br>1 day $NH_3$ | 1 day PBS<br>1 day $NH_3$ | 6 days PBS<br>1 day $NH_3$ | 7 days PBS | 1 day $NH_3$ |
|---|---|---|---|---|---|---|
| **Casted** | 24.5 ± 5.4 | 13.8 ± 3.5 | 20.4 ± 4.0 | 32.8 ± 10.4 | 26.18 ± 12.5 | 16.6 ± 4.1 |
| 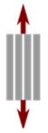 | 25.0 ± 9.4 | 16.3 ± 1.5 | 25.1 ± 9.6 | 17.3 ± 9.9 | 10.8 ± 1.6 | 16.0 ± 4.2 |
| 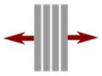 | 8.5 ± 1.9 | 18.7 ± 1.8 | 16.0 ± 7.5 | 13.0 ± 2.9 | 6.4 ± 0.9 | 13.0 ± 3.9 |

**Figure S4.** Young's modulus calculated on the small deformations for all types of gels.

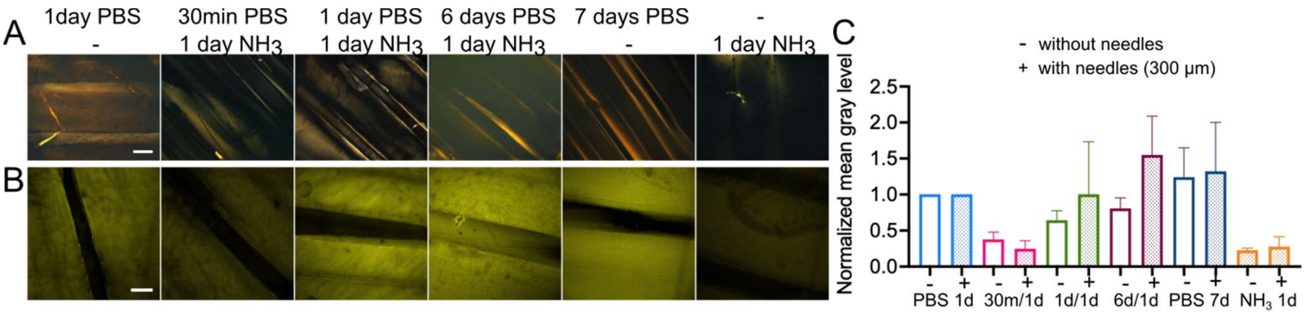


**Figure S5.** Generation of large channels within printed hydrogels without alteration of collagen anisotropy. (A) Polarized light microscopy on 3D-printed hydrogel slices with the presence of needles. (scale bar 500 µm). (B) Second-harmonic generation imaging of slices of printed gels with needles (scale bar 250 µm). (C) SHG intensity of printed hydrogels with needles (+) compared to those without needles (-).

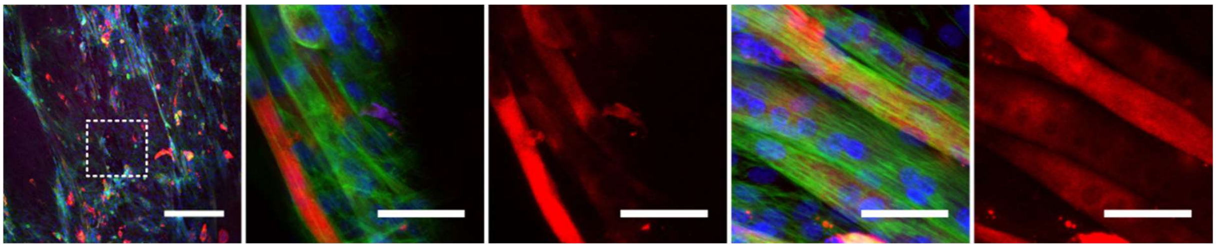

**Figure S6.** C2C12 seeded into printed gels for 7 days. Left: Longitudinal view of large pores (scale bar 100 µm). Green: actin, blue: nucleus, red: MF20. Zoom on myotubes alignment and myosin staining (scale bar 50 µm).
